# Towards an Eye-Brain-Computer Interface: Combining Gaze with the Stimulus-Preceding Negativity for Target Selections in XR

**DOI:** 10.1101/2024.03.13.584609

**Authors:** G. S. Rajshekar Reddy, Michael J. Proulx, Leanne Hirshfield, Anthony J. Ries

## Abstract

Gaze-assisted interaction techniques enable intuitive selections without requiring manual pointing but can result in unintended selections, known as Midas touch. A confirmation trigger eliminates this issue but requires additional physical and conscious user effort. Brain-computer interfaces (BCIs), particularly passive BCIs harnessing anticipatory potentials such as the Stimulus-Preceding Negativity (SPN) - evoked when users anticipate a forthcoming stimulus - present an effortless implicit solution for selection confirmation. Within a VR context, our research uniquely demonstrates that SPN has the potential to decode intent towards the visually focused target. We reinforce the scientific understanding of its mechanism by addressing a confounding factor - we demonstrate that the SPN is driven by the user’s intent to select the target, not by the stimulus feedback itself. Furthermore, we examine the effect of familiarly placed targets, finding that SPN may be evoked quicker as users acclimatize to target locations; a key insight for everyday BCIs.

**CCS CONCEPTS:**

- **Human-centered computing** → **Virtual reality**; **Mixed / augmented reality**; *Accessibility technologies*; **Interaction techniques**.

**ACM Reference Format:** G. S. Rajshekar Reddy, Michael J. Proulx, Leanne Hirshfield, and Anthony J. Ries. 2024. Towards an Eye-Brain-Computer Interface: Combining Gaze with the Stimulus-Preceding Negativity for Target Selections in XR. In *Proceedings of the CHI Conference on Human Factors in Computing Systems (CHI ‘24), May 11–16, 2024, Honolulu, HI, USA*. ACM, New York, NY, USA, 17 pages. https://doi.org/10.1145/3613904.3641925

## 1 INTRODUCTION

Gaze-based interactions have emerged as a powerful and intuitive method for users to interact with Extended Reality (XR) environments. This surge in interest can be attributed not only to the growing number of Head-Mounted Displays (HMDs) incorporating eye-tracking systems [93], but also to the efficiency, speed, and reduced physical effort that gaze interactions offer [105, 110]. Essentially, these interactions eliminate the need for the pointing step in *point-and-select* interactions. Numerous studies have investigated how to harness natural eye movements for direct input, such as object selection [2, 14, 71, 74, 121], with one of the most common methods being the Dwell technique [25, 30, 43, 76]. This method mandates users to fixate on a target for a predetermined duration to confirm selection, distinguishing between passive viewing and active engagement. However, the Dwell technique is not without its drawbacks, as it can lead to the Midas touch issue [43], where unintentional selections are mistakenly registered by the system.

To mitigate the Midas touch, HCI researchers have proposed the use of an external trigger as a form of manual input [128]. This solution has been implemented in a variety of ways from voluntary blinks [71] to simply pressing a button on a physical controller [29] (see Section 2.1). These techniques improve selection accuracy, however, they require physical exertion hence potentially introducing fatigue over extended use [45]. Furthermore, these actions diminish the instinctive nature of gaze interactions and can break the immersive experience, as users are forced to shift their attention to voluntarily execute a motor task. The pursuit of a more effortless and passive approach to addressing Midas touch in gaze-based interactions continues to be a significant challenge, especially for XR experiences.

Brain-computer Interfaces (BCIs) have the potential to overcome these limitations by offering a natural and accessible interaction technique for these spatial environments [39]. However, prevalent BCI paradigms, including the P300 Event Related Potential (ERP), Steady State Visually Evoked Potential (SSVEP), and Motor Imagery (MI)-based BCIs are fraught with issues such as high user-workloads, fatigue, and slow interaction times that limit their widespread adoption [1], especially for healthy users. Typically, these BCIs are categorized as *active*, where users need to consciously imagine muscle movement (MI), or *reactive*, where reactions such as phase-locked spectral activity are evoked by stimuli such as SSVEP’s flashing lights [3]. In contrast, *passive* BCIs have emerged as a compelling avenue, where users do not deliberately control or manipulate their thoughts nor does the stimuli need to be altered to evoke a specific brainwave response, facilitating smoother interactions and reducing user cognitive load [126]. Passive BCI paradigms tapping into slow negative waves, specifically the Stimulus-Preceding Negativity (SPN) and the Contingent Negative Variation (CNV), have gained attention only in recent literature [41, 94, 104, 123, 129], though they were proposed back in 1966 for ‘the direct cerebral control of machines’ [118].

The SPN is a purely nonmotor negative potential that arises when a user expects a forthcoming stimulus, often as a consequence of a prior action [8]. It results from the summation of numerous postsynaptic potentials in the brain regions that will be engaged in processing the upcoming stimulus. Its functional importance is believed to be related to preparatory activity or the suppression of neural activity, aimed at accelerating neural processing upon receiving the stimulus [8]. This negative potential has the capability to serve as a passive confirmation mechanism (to effectively address the Midas touch), and we investigate that capability in this work. Moreover, SPN allows for an unobtrusive and implicit integration of user intent detection into the interaction process, thus making it *seamless*.

To understand how an SPN can act as a confirmation trigger, let’s consider a user navigating an XR music interface. As the user gazes at the *play* button, they anticipate a stimulus feedback (an audio confirmation or a visual cue such as a change in UI) that will happen upon selection. The SPN is elicited due to their anticipation of the stimulus that will be provided upon selection. The system, sensing the change in brainwaves via Electroencephalogram (EEG), can automatically activate the *play* function. Such an interaction, where a user’s anticipation of a response becomes the trigger, enables a more fluid interaction. The user no longer has to make a deliberate action, but instead, their inherent neural signatures based on experience and intent becomes the primary mode of interaction. Thus, these negative slow-wave potentials serve as a *passive* mental switch and have the potential to address inadvertent gaze selections as users do not expect feedback when not selecting a target. More importantly, this *passive* mechanism sidesteps the need for users to consciously engage in the selection process, eliminating the selection step as well in *point-and-select* interactions.

In this study, we delve deeper into the dynamics of SPN in the context of XR, by examining its presence within an immersive three-dimensional VR environment (Figure 1) and exploring its viability as a solution to the challenges presented by the Midas touch issue in gaze-based dwell interactions. We also address certain potential confounds, unclarified from prior research, thereby establishing SPN’s link to user anticipation, and examine the effect of UI familiarity on its manifestation.

**Figure 1:**
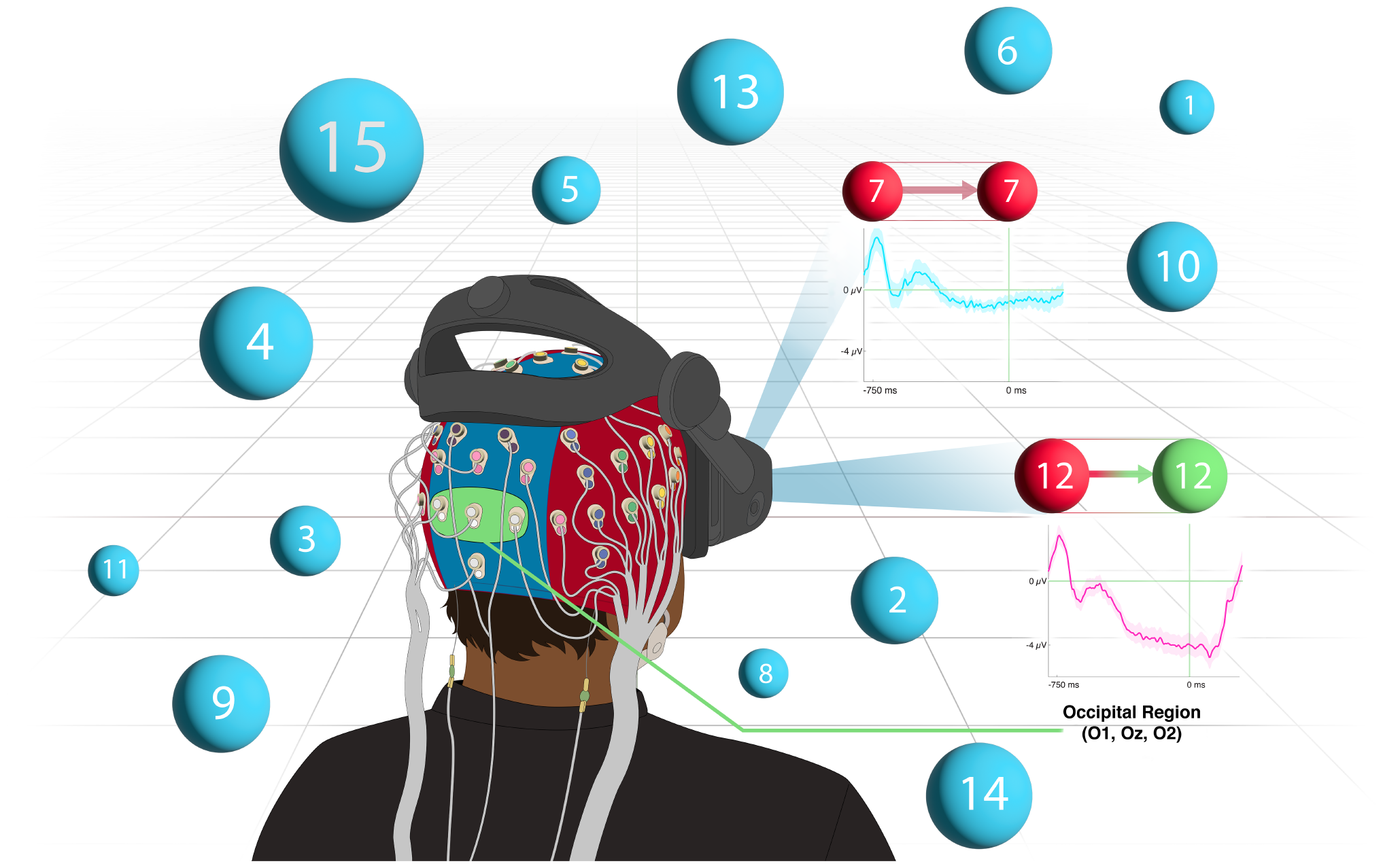
**An illustration of our approach to address the Midas touch issue in gaze selections for immersive environments. When the user merely observes target 7, no Stimulus-Preceding Negativity (SPN) is evident from the occipital region’s electroencephalogram activity. When intending to select target 12 and expecting a selection feedback, an SPN is evoked, serving as an implicit selection trigger**.

### Terminology

Previous neuroscience research [104, 129] that explored SPN in addressing the Midas touch, differentiated active engagement and passive viewing in Dwell-based control using terms such as *Intentional vs Non-intentional fixations* and *Intentional vs Spontaneous selections*. However, these distinctions may be misleading within the HCI field, as all gaze activities are fundamentally intentional. Therefore, in our study, we adopt the terms *intent-to-select* interactions for the explicit intent to interact with or select a target and *intent-to-observe* interactions for implicit or passive engagement, such as watching or viewing, without intending to interact with it. Our choice of terminology more accurately and precisely captures the distinction in the user’s intention and the intent behind their gaze activity.

## 2 RELATED WORK

This work builds on insights from past research that have addressed the Midas-Touch issue in gaze-based *point-and-select* interactions.

### 2.1 Gaze-based Interactions

The Dwell technique has been the most common gaze interaction technique [24, 25, 30, 43, 75, 76] in HCI literature. Prior works like Jacob [43] used Dwell for selections (who also first noticed the Midas touch) and other works like Starker and Bolt [109] leveraged Dwell to make the system provide more information after a fixed period, sort of an interest metric. Past research has also explored optimal parameters for the Dwell technique, such as minimum target size [120] or dwell threshold [75, 98, 116], finding that it is heavily dependent on the task, context, and cost of errors. [90].

Researchers have studied gaze as both an individual and a multimodal system, where other modalities usually confirm the user’s gaze target [107, 108], addressing the Midas touch. Hence, it is a two-step procedure where the user’s gaze acts as a cursor and the selection is triggered by an additional modality. Various modalities have been investigated, including eye blinks [71], speech [63, 96, 122], head movements [115, 121], handheld devices [29, 64], tongue-based interfaces [34], body movements [57], and hand gestures such as the pinch gesture [64, 82, 99]. Additional gestures, such as gaze-hand alignment [74] confirm target selection either by aligning a ray that emanates from the user’s hand with their gaze (Gaze&Handray) or aligning the user’s finger with their gaze (Gaze&Finger) [117]. However, gestural interfaces can lead to physical fatigue [45], making them difficult to use for extended periods. Gaze interactions may also cause eye fatigue or strain [97], but the direct measurement of their impact is not yet known, as they are assessed using devices that themselves induce fatigue symptoms [37]. Moreover, research has found less eye strain when gaze was used only for pointing, rather than being used for both pointing and selection [68].

Research has also explored using natural eye movements as confirmation triggers, such as voluntary blinks [71]. In contrast, some studies have explored half-blinks [65], removing the need to differentiate between spontaneous and voluntary blinks. Smooth Pursuits [114], offer another approach that bypasses the need for precise gaze tracking or a prior eye-calibration step, and this technique has found applications in XR environments [28, 52]. Vergence-matching [2, 112] is a method that gauges the simultaneous opposing movement of both eyes, correlated with subtle changes in a target’s depth. If the vergence movement corresponds to a specific target’s depth alteration, that target is identified as the user’s point of focus. Though this approach has proven effective in selecting small targets [106], it is often perceived as uncomfortable, difficult to execute consistently, and diverts the user’s attention [56].

Model predictions using head and eye endpoint distributions were explored by Wei et al. [121], taking into account the distributions of the head and eye positions/orientations during target selection. They developed a robust target prediction model that surpassed the baseline. Other works like Bednarik et al. [5] have explored predicting intents using gaze metrics alone such as saccade and fixation durations, saccade amplitudes, and pupillary responses. They achieved an accuracy of 76% with Area Under Curve (AUC) around 80%. David-John et al. [14] similarly explored intent prediction in VR, using gaze features such as the K coefficient [61], gaze velocity, and so on; predicting intent significantly above chance with the the Receiver Operating Characteristic(ROC)-AUC scores showing a maximum of 0.92 and a minimum of 0.71.

### 2.2 Gaze-assisted Neural Interactions

Gaze signals, when combined with BCIs, give rise to hybrid BCIs (hBCIs). This fusion is commonly adopted to increase the range of control commands, enhance classification accuracy, or reduce the intent detection time [39, 91]. In studies conducted between 2009 to 2016, 52% of them investigated this merger of gaze and neural signals [39], including gaze captured from both Electrooculogram (EOG) and an eye-tracker. EOG signals, known for their temporal precision, are primarily used to reduce ocular artifacts leading to lesser false positives in classification [4, 40, 81, 113]. Gaze signals have also been used to improve control commands such as the control of a wheel-chair [55], where the classifier achieved an accuracy of 97.2%. Kalika et al. [47] combined gaze with EEG in a P300-based speller system. The integrated system surpassed the EEG-only setup in accuracy and bit rate.

Eye blinks have also been incorporated with both MI and P300-based BCI paradigms for wheelchair control [119]. Pfurtscheller et al. [91] examined an MI-based hBCI solution to the Midas touch problem using eight subjects in a search-and-select task. The hybrid approach (EEG + Gaze) proved significantly more accurate in selections compared to the Dwell technique implemented using gaze alone. Nonetheless, the hBCI was the slowest activation method, while demanding conscious effort from the user. Other works have investigated MI and gaze-based BCIs, achieving higher accuracy, yet voluntarily imagining motor movements was challenging for users [20].

A notable application by Kim et al. [54] showcased a unique combination: gaze signals from an eye-tracker combined with the SSVEP paradigm to direct a turtle’s movement. Directional commands (e.g., turning left or right) were controlled by SSVEP, whereas *reset* and *idle* commands were activated by simply opening or closing the eyes. McCullagh et al. [80] also examined the SSVEP with the Dwell technique, finding users lacking technical proficiency, encountered difficulties navigating the interface. In a related study, Putze et al. [95] combined gaze with SSVEP for AR-based smart home management, yielding better accuracy than SSVEP by itself. Nonetheless, the flashing visual stimuli of SSVEP targets disrupted users’ cognitive flow.

### 2.3 Passive Hybrid BCIs

A passive hBCI combines signals from multiple modalities, and derives its output from implicit brain activity without demanding active user participation, with the goal of enhancing interaction techniques by making them more seamless and effortless. Prior works have explored such interfaces like Zander et al. [127], who used error-related potentials (ErrPs) to guide a two-dimensional cursor to its desired target based on implicit user states from the pre-frontal cortex. If the cursor’s position mismatched the user’s expectation, an ErrP was triggered, with its peak amplitude correlating with the error magnitude. In a similar vein, Krol et al. [62] designed a Tetris game where gaze and ErrPs were the primary controls: Gaze determined block movements while factors like the player’s relaxed state influenced game speed, and ErrPs eliminated undesired blocks. Others have also employed ErrPs to rectify inadvertent Dwell technique selections [46].

Research efforts on passive BCIs has also sought to classify user intent. For instance, Sharma et al. [102] achieved a 97.89% accuracy in discerning intent during free-viewing, target searching, and target presentation conditions, utilizing EEG features such as power spectral intensity and detrended fluctuation analysis. However, given the task’s emphasis on visual searches, it’s likely that the primary EEG components were the P300 wave or reward potentials [3, 32]. This suggests that the model’s accuracy might differ when selecting familiar targets, as is the case for everyday UI interactions. Previous studies have also investigated the classification of user intent to interact with virtual objects by analyzing EEG and Electromyography signals, achieving detection accuracies of pre-movement states up to 69% [84]. While not directly stated, this research likely leveraged the CNV wave, an anticipatory potential akin to SPN but linked to motor activity [8]. Kato et al. [49] also used the CNV to develop a switch for activating or deactivating a BCI system. In other works [123], the CNV was used in conjunction with a P300-based BCI to improve its accuracy from 85% to 92.4% in identifying visual targets. While the CNV appears promising as a passive mental ‘click’, its reliance on imagined motor activity can distract users from their primary task.

Conversely, the SPN is tied to nonmotor activity [8]. An early study by Protzak et al. [94] used this slow negative wave, achieving a mean accuracy of 81% when distinguishing between intent-to-select and intent-to-observe interactions. This was attributed to pronounced negativity in the parietal region of interest (ROI) during intent-to-select interactions. The study involved target selections on a two-dimensional screen using a 1000ms Dwell threshold. The SPN in this study was associated with the stimuli indicating a successful selection, i.e., the task proceeding to the next trial, rather than being an explicit, separate visual or auditory stimulus. However, the study focused on visual searches where the prominent P300 component [3, 32] could influence classification results, potentially affecting intentions towards familiar targets. Moreover, the study was limited by the lack of tests for statistical significance and absence of details on ocular artifact corrections.

Shishkin et al. [104] demonstrated that the SPN, in the parietooccipital ROI, can distinguish intent-to-select from intent-to-observe fixations within 300ms segments. In their study, eight participants engaged in a gaze-controlled game, using the Dwell technique to select and maneuver game pieces. The authors explored both 500ms and 1000ms dwell thresholds, which were found not to influence the SPN. Visual feedback was provided for intent-to-select interactions only, likely to induce the SPN wave. While EOG was recorded, it was not used for ocular artifact correction. Not correcting for these artifacts can increase false positives in classification [4, 40, 81, 113]. Instead, the authors analyzed windows believed to be free from such contamination.

In a later study, Zhao et al. [129] corroborated the SPN’s presence in voluntary selections of moving objects and successfully distinguished between intent-to-select and intent-to-observe smooth pursuits in 8 participants among 20. They employed a two-dimensional task where participants explicitly selected 15 targets sequentially, while intent-to-observe interactions involved a counting task. Notably, heightened negativity was recorded at the Oz and POz electrodes during intent-to-select pursuits compared to intent-to-observe ones. Only intent-to-select interactions received visual feedback, where the target turned green upon a successful selection.

## 3 NOVELTY, HYPOTHESES, AND CONTRIBUTIONS

Prior work [94, 104, 129] has demonstrated the SPN’s ability to distinguish between an intent-to-select and an intent-to-observe interaction, indicating its potential in resolving the Midas touch problem in gaze-based dwell interactions, via a more seamless or unobtrusive approach. However, having used two-dimensional stimuli on computer monitors, such studies are often constrained to a small visual field, utilizing screen-based eye trackers with restricted head movements [104, 129]. They offer limited ecological validity for understanding human behavior in more naturalistic 3D environments.

In distinguishing our research’s use of XR environments, it’s imperative to consider its unique aspects. XR’s immersive nature, coupled with its depth and spatial dynamics, offers a complex interaction space as illustrated in Figure 1. This 3D environment influences cognitive processes and neural responses, potentially in how users anticipate and react to stimuli. Prior work found that 3D immersive environments, in comparison to 2D, decreases the alpha band power and amplifies cortical networks in the parietal and occipital lobes [51, 70, 124], due to the enhanced visual attention and perception. Li et al. [67] adds that VR intensifies visuospatial sensory inputs, leading to increased corticothalamic connectivity [103]. This enhanced connectivity, in turn, leads to a more rapid cortical spike timing, as evidenced by the quicker onset of the P300 component [10]; which is also linked to visual stimuli like the SPN.

Moreover, much of our current knowledge derived from 2D stimuli studies on monitors, does not effectively apply to XR environments. For instance, research indicates that VR environments can reduce working memory reliance in object search and copying tasks, compared to traditional laboratory settings [23]. They also found that as locomotive demands increase, such as when gaze moves to more peripheral locations, the dependence on working memory increases. Lastly, the integration of head and eye movements in XR enhances the realism, user engagement, and provides a richer context for studying anticipatory neural responses like the SPN.

We also introduce an additional control condition where feedback is presented to the user even when they are not actively trying to select the target, in contrast to prior work [104, 129] that did not have this controlled between conditions. This condition seeks to confirm that the SPN is intrinsically tied to user’s implicit *anticipation* of selection feedback, regardless of feedback presence. It is worth noting here that, to the best of our knowledge we are the first to address this confound.

Additionally, we contrasted the ERPs from intent-to-select interactions between instances when participants were familiar with target locations and when they were not, to determine whether an SPN manifests more rapidly when targets are familiar. Prior work has shown that memory guided attention can trigger temporal predictions leading to enhanced performance and neural activity at the expected location [12, 86]. We particularly investigated this because in a BCI used for everyday tasks, users are typically well-acquainted with the UI they engage with. Understanding how familiarity with the UI influences anticipatory responses is vital for optimizing BCI systems for everyday use. Building on the foundation laid by prior research, we set forth specific hypotheses:

- **H1:** We hypothesize the SPN will manifest in an intent-to-select interaction but not in an intent-to-observe interaction for an immersive environment.
- **H2:** We hypothesize that an SPN is driven by the user’s intention, or anticipation of the stimulus, and not by the stimulus presence itself.
- **H3:** As users acclimatize to the target locations, we hypothesize the SPN’s quicker manifestation.

## 4 USER STUDY

Our study aims to ascertain whether the SPN wave, a type of slow negative potential, appears during an intent-to-select interaction in an XR environment, a key preliminary investigation in addressing the Midas touch issue. To explore this, we devised a VR task involving three-dimensional object selection using the Dwell technique.

### 4.1 Participants

The University of Colorado recruited 22 participants through online bulletins, but rejected 5 due to technical issues, difficulties with the counting task, or the eye-tracker’s failure to accurately track gaze. 17 participants (9 male and 8 female), averaging 28.8 years (SD = 10.7) are analyzed in this work. One participant had moderate experience with virtual and augmented reality, 10 participants had minimal experience, and 6 had no experience; 5 participants had experience with gaze-based control. Participants who required corrective eye-wear were excluded due to gaze tracking concerns, although clear contact lenses were allowed and 4 participants wore them; 9 participants had naturally normal vision and 8 had corrected-to-normal vision. All participants reported free of any neurological disorders. They were paid 15 USD per hour for the two-hour study.

### 4.2 Apparatus

The experimental setup, as seen in Figure 2, was comprised of an (i) HP Reverb G2 Omnicept Edition VR headset with an integrated Tobii eye-tracker ^1^, and (ii) a 64-channel BioSemi ActiveTwo EEG system^2^ with four additional EOG electrodes and two reference electrodes that were recorded on the same amplifier. 64 EEG electrodes were placed in a 10-10 montage across the scalp [87]. The reference electrodes were placed on the left and right mastoids, and the EOG on the outer canthus and below the orbital fossa of each eye. We made four slits in the HMD’s face cushion to reduce recording interference and pressure points on the EOG electrodes. All electrode impedances were kept below 50 kΩ and referenced to the Common Mode Sense active electrode during the recording. EEG and EOG data were recorded at 512 Hz. The Tobii eye-tracker, sampling at 120Hz, identified the user’s gaze point in real time, while also logging pupillometry at the same rate.

**Figure 2:**
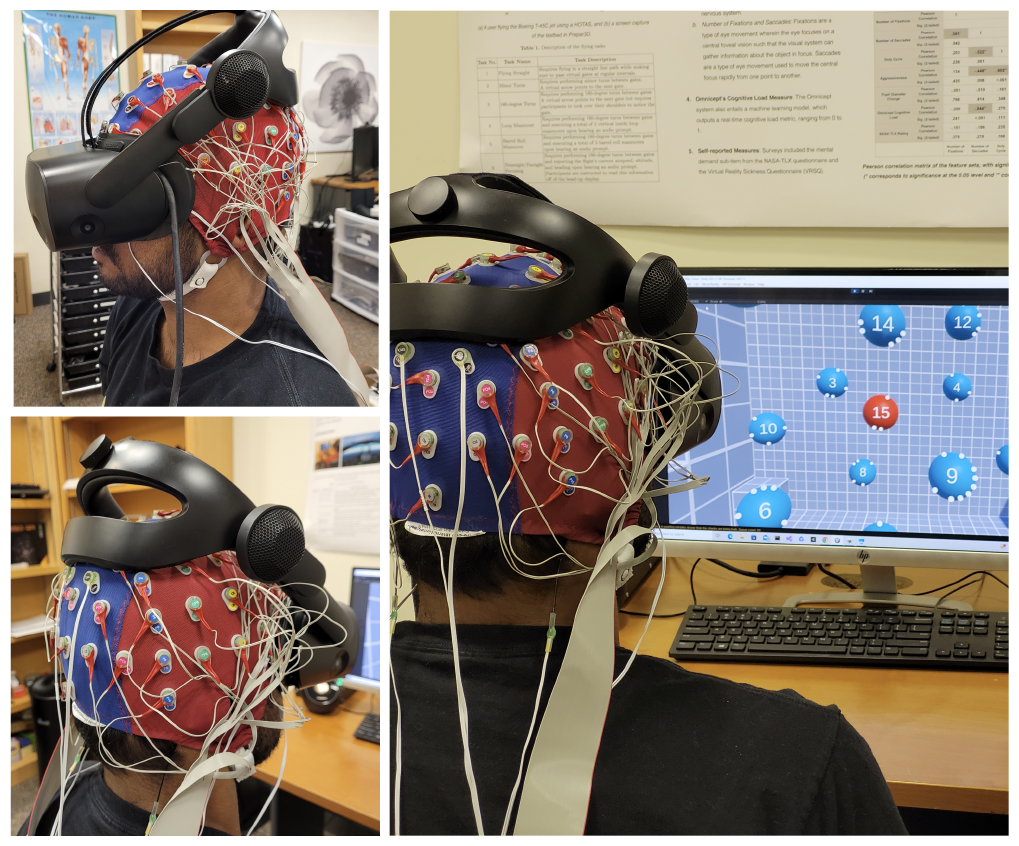
**A user selects targets while EEG, EOG, and eye-trackers capture their physiological responses. The monitor displays the user’s VR view, with the red target indicating their current visual focus**.

The VR headset was tethered to a computer with an Intel i7-9700K @ 3.6GHz CPU, RAM of 16GB, and an NVIDIA GeForce RTX 2080 Super GPU. Our tasks, which were developed and run on Unity^3^, maintained a 90fps frame rate, fully leveraging the HMD’s display capacity. The HMD used inside-out tracking for six-degrees-of-freedom tracking, had a resolution of 2160 × 2160 per eye, and a 114^°^ field of view. A separate computer was connected to the BioSemi amplifier, on which all data was logged. The EEG, EOG, eye-tracker, and an experimental marker stream (which highlighted interaction events and other key moments in our task) were synchronized and logged using the Lab Streaming Layer^4^, facilitated by wired LAN connections between the systems.

### 4.3 Task

Participants engaged in VR-based three-dimensional object selection tasks modeled after Zhao et al. [129], though the objects remained static in our study. The objects comprised of 15 targets, i.e., numbered blue spheres spread throughout a three-dimensional space as depicted in the left image of Figure 3. The nearest target occupied a visual angle of 19.6°, and the farthest target had a visual angle of 7.8°. The targets were distributed across an area that spanned a horizontal visual angle of 60.4° and a vertical visual angle of 47°.

**Figure 3:**
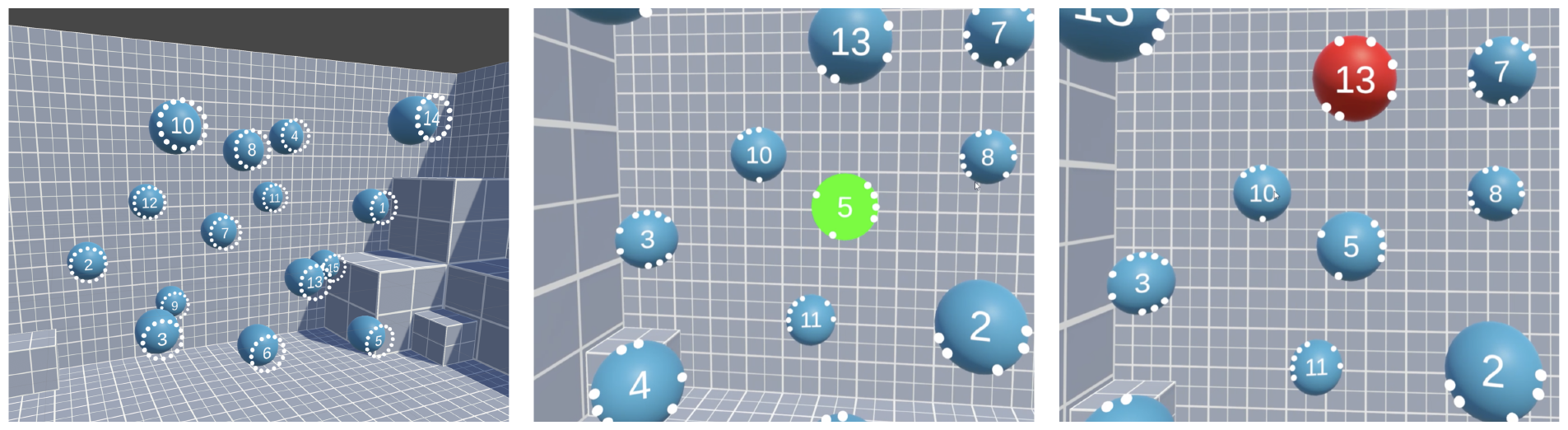
**(Left) 15 numbered targets dispersed across a three-dimensional space in the Unity game engine. (Middle) A screen capture of the user’s view depicting the target’s bright green stimuli in the Intent-to-select and Intent-to-observe with feedback conditions. (Right) A screen capture of the user’s view depicting the Intent-to-observe without feedback condition where the target remained red, indicating the user’s current visual focus**.

#### Dwell-based selection mechanism

The HMD’s eye-tracker provided real-time coordinates of the gaze origin and the normalized gaze direction. Using Unity’s *Raycast* function, we projected a virtual ray from the gaze origin in the direction of this normalized vector. The point at which this ray intersected an object in the environment indicated the user’s current visual focus. When the user’s gaze landed on a target, the ray intersected the target’s collider, changing its color from blue to red, as seen in the right image of Figure 3.

Prior research found the presence of the SPN at dwell thresholds of both 1000ms and 500ms [104]. We chose a 750ms threshold for our study as a balanced approach, considering the novel challenge of detecting an SPN in immersive environments. The dwell countdown started when the user’s gaze first landed on the target (or when it turned red). If the user’s gaze remained fixed on that target for the full 750ms duration, the target illuminated in bright green (as seen in the middle image of Figure 3) accompanied by the audio cue, signaling a successful selection. It is this bright green visual stimuli that we expect the user to anticipate in intent-to-select interactions. The target turned back to blue when the user’s gaze shifted away from it.

The tasks involved interactions across three conditions: (i) Intent-to-select, (ii) Intent-to-observe without feedback, and (iii) Intent-to-observe with feedback.

#### 4.3.1 Intent-to-select Interactions

In this condition, participants were required to select the 15 targets based on the Dwell mechanism, with targets turning green and accompanied by a short audio cue to indicate a successful selection. The Intent-to-select condition was split into two separate tasks: selecting targets in ascending (Intent-to-select ascending) and in descending order (Intent-to-select descending) — to ensure a thorough mix with the Intent-to-observe conditions.

#### 4.3.2 Intent-to-observe Interactions

For intent-to-observe interactions, which served as the control, participants engaged in a counting task without intending to select the targets. These conditions depict instances where inadvertent dwell selections can happen when users are passively observing the target. Each sphere had varying surrounding dots and participants were instructed to identify five spheres with a specific number of dots. Upon identifying them, they were instructed to add up the central numbers displayed on the spheres. The resulting sum was then entered into a virtual keypad in VR using a controller. The intent-to-observe interactions were split into two conditions:

##### Intent-to-observe without feedback

In this condition, the Dwell selection mechanism operates similarly in the background. However, unlike the Intent-to-select condition, the target does not turn green post the Dwell threshold because the user is not “selecting” the target. Nevertheless, an intent-to-observe interaction trigger is logged once the timer ends.

##### Intent-to-observe with feedback

If the stimulus is not provided during intent-to-observe interactions, is the missing SPN attributable to the lack of stimulus or the lack of the user’s intention to select? To address this ambiguity, we introduced an extra control condition. In this condition, the target turns green and is accompanied by the audio cue despite the user not actively selecting the target. We hypothesize **(H2)** an SPN’s absence in this condition.

To summarize, the study consisted of three conditions (Intent-to-select, Intent-to-observe without feedback, and Intent-to-observe with feedback), presented across four tasks: (i) Intent-to-select ascending, (ii) Intent-to-select descending, (iii) Intent-to-observe without feedback, and (iv) Intent-to-observe with feedback.

### 4.4 Experimental Procedure

We employed a within-subjects design, with participants completing three blocks, each containing five phases as shown in Figure 4. In each phase, the four tasks (depicted by the blue boxes) were randomized. Participants performed a total of 450 intent-to-select interactions across all three blocks. Each block consisted of 150 interactions, further broken down into 30 interactions for each phase, with each task comprising 15 interactions. For intent-to-observe interactions, there was no fixed number of interactions as the task was designed to be exploratory. Nonetheless, each task required participants to identify at least 5 targets marked with the specified number of dots, resulting in a minimum of 10 interactions for each phase. This translates to at least 50 interactions in each block and a total of 150 interactions minimum across all three blocks. The actual number of interactions varied from task to task for every participant.

**Figure 4:**
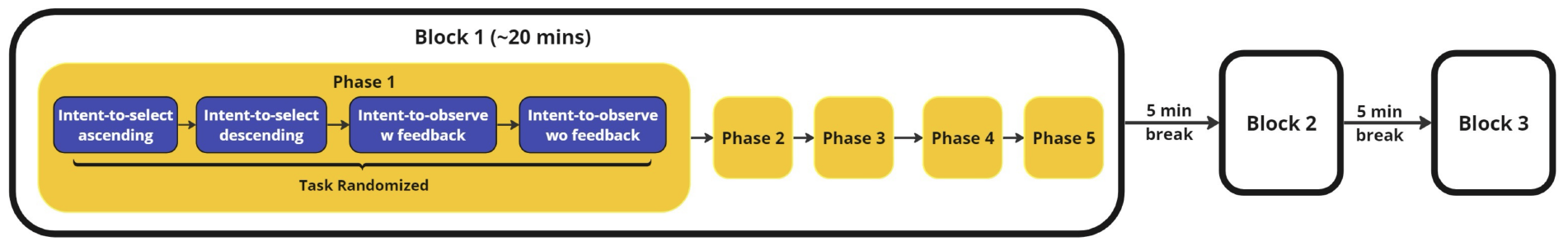
**Experiment timeline depicting the sequence of tasks. Each block comprises five phases with the four tasks’ (Intent-to-select ascending, Intent-to-select descending, Intent-to-observe with feedback, Intent-to-observe without feedback) order randomized within each phase**.

At the start of every intent-to-observe interaction task, the number of dots to find, the number of dots present on each target, and the position of the dots on each target were randomized, with the number of dots to find falling within a range of 5 to 8. To aid familiarity with target locations, the target numbers remained in the same spatial locations within each block (phases 1-5). This was done to specifically probe **H3**. However, these numbers were randomized at the start of each block.

Upon arriving at the lab, participants signed consent forms per the University’s IRB (IRB# 23-0156) and completed a demographics questionnaire. Next, the EOG, reference, and EEG electrodes were attached to the participants, with electrode gel applied to ensure optimal signal conductance. Participants received task instructions, with specific guidelines such as making slow head movements. They were advised to approach the tasks at a comfortable pace and to focus exclusively on the tasks. They were also cautioned against speaking during the study and were reminded to maintain relaxed facial muscles to prevent interference with EEG signals. Before starting the tasks, participants wore the HMD and underwent an 8-point native eye-calibration.

Before the main task, participants completed a tutorial encompassing one phase of all four tasks. They then proceeded to complete the three blocks, each lasting approximately 20 minutes and each followed by a 5-minute break where they could remove the HMD. The eye-tracker was recalibrated at the beginning of each block and any time the participant put the HMD back on after removal.

### 4.5 Eye/EEG Data Processing

Data processing and analysis were conducted offline through custom MATLAB scripts leveraging the EEGLAB toolbox [16], along with ERPLAB [69] and the EYE-EEG plugins [19]. The eye-tracking data was used for the detection of eye events. First, blinks were identified by recognizing typical gaps in gaze behavior ranging from 50 to 700ms [38]. Any gaps shorter than 50ms were interpolated and those larger than 700ms were deemed data dropouts. Subsequently, using a velocity-based algorithm [26, 27], adapted from the EYE-EEG plugin [19], we identified saccade and fixation events. Specifically, a velocity factor of six (or six times the median velocity per subject) combined with a minimum saccade duration of 12ms was applied. Fixation events were recognized as eye data with velocities below this velocity factor. To remove biologically implausible eye movements and outliers obtained from the previous velocity factor classification, we focused on fixations that lasted between 100 to 2000ms and saccades meeting the following criteria: (i) a 12 to 100ms duration, (ii) a 1 to 30° amplitude, and (iii) a velocity ranging from 30 to 1000°/sec. We then merged the eye-tracker data with the EEG data (upsampling the eye data to match EEG), including the identified blink, saccade, and fixation events.

#### 4.5.1 Correcting ocular artifacts

In complex visual search tasks, ocular artifacts can be introduced due to the movement of the eyeballs, eyelids, and extraocular muscles that contaminate the neural activity [92]. To isolate EEG recordings associated with these artifacts, we applied an ICA to the EEG and EOG data. This was done using the extended Infomax algorithm [66] in line with the OPTICAT methodology [18]. The OPTICAT method is particularly effective in removing eye muscle-generated saccadic spike potentials [18]. It employs a more liberal filter settings to ensure retention of higher frequency activity in the training data, overemphasizes saccade onset events, and automates the process of identifying and discarding inappropriate ICs.

For every participant, we initially re-sampled the EEG data from 512 Hz to 256 Hz across all 70 EEG, EOG, and mastoid channels and then re-referenced the data using the average of the left and right mastoids, in line with previous studies [17, 33, 50, 85, 89]. To detect bad channels, we applied a voltage threshold of four standard deviations, using the *pop_rejchan* function in EEGLAB. These bad channels were then spherically interpolated, using the EEGLAB’s *pop_interp* function. Finally, we applied a second-order Butterworth filter with a 2 to 100 Hz bandpass filter, optimal for our task [18].

Before applying ICA to the EEG data, we overweighted saccade events by considering EEG intervals 20ms before and 10ms after saccade onsets, resulting in twice the original event data. After running ICA, we used an automatic IC flagging/rejection process to classify activity related to ocular artifacts based on a procedure from Plöchl et al. [92] implemented in the EYE-EEG toolbox (function: *pop_eyetrackerica*). We computed variance ratios between saccade epochs and fixation epochs (i.e., variancesaccade/variancefixation) using a saccade window starting 11.7ms (or 3-samples) before saccade onset until the saccade offset and fixation windows defined as from fixation onset to 11.7ms before a saccade. If the variance ratio exceeded 1.1, it was flagged as an eye-artifact related IC [92]. On average, 4.2 ICs (SD = 1.26) were marked for rejection across all participants. The benefit of this approach is that it uses an objective criteria to identify which ICs covary with saccade and fixation related activity measured by an eye-tracker.

### 4.6 Extracting ERPs

The preproccessing for the ERPs followed an approach akin to the ICA preprocessing. We resampled the EEG signals from 512 Hz to 256 Hz, eliminated 60 Hz line noise using the Zapline-plus extension for EEGLAB [15, 58], and re-referenced the data using the average of the left and right mastoids. Bad channels were interpolated as mentioned earlier, followed by application of a 4th order 0.1 to 40 Hz Butterworth bandpass filter. ICs marked for rejection during ICA decomposion were removed and the remaining ICs were back-projected to this new pre-processed file. In this way, the ocular artifacts could be substantially reduced in our EEG signal. We used ERPLAB to epoch the data, extracting ERPs time-locked to the selection / interaction event within a -2000ms to 1500ms window.

For intent-to-select interactions, only the events pertaining to correct selections (sequentially triggered) were used, and inadvertent selections were discarded. Moreover, intent-to-select interactions were reduced to the mean of the ascending and descending tasks for further analysis. For intent-to-observe interactions, all trials that led to the interaction trigger (750ms) are used, given that the accuracy of the counting task is not pertinent to our analysis.

The ERPs were baseline corrected from -1000 to -850ms as we were primarily interested in the -750 to 0ms window where an SPN is to be expected. We avoided the -850 to -750ms range due to the fixation-related Lambda response resulting from the first fixation on the target stimulus. ERP epochs with artifacts were detected and excluded using a 100μV sample-to-sample voltage threshold, resulting in roughly 2.6% rejection rate per condition across all participants. The average number of trials used for the grand average ERP for each task is shown in Table 1. We focused our analysis on the occipitoparietal ROI (O1, Oz, O2, Iz, PO7, PO3, POz, PO4, PO8, P5, P6, P7, P8, P9, and P10 electrodes as depicted in Figure 5), in line with prior findings highlighting the SPN’s prominence in these regions [94, 104, 129].

**Table 1:** Number of ERPs extracted for each task, averaged across 17 participants.

| Task | # (%) Accepted | # (%) Rejected |
| --- | --- | --- |
| Intent-to-select ascending | 3645 (98%) | 75 (2%) |
| Intent-to-select descending | 3635 (97.7%) | 84 (2.3%) |
| Intent-to-observe without feedback | 3824 (97%) | 118 (3%) |
| Intent-to-observe with feedback | 3594 (96.8%) | 120 (3.2%) |

**Figure 5:**
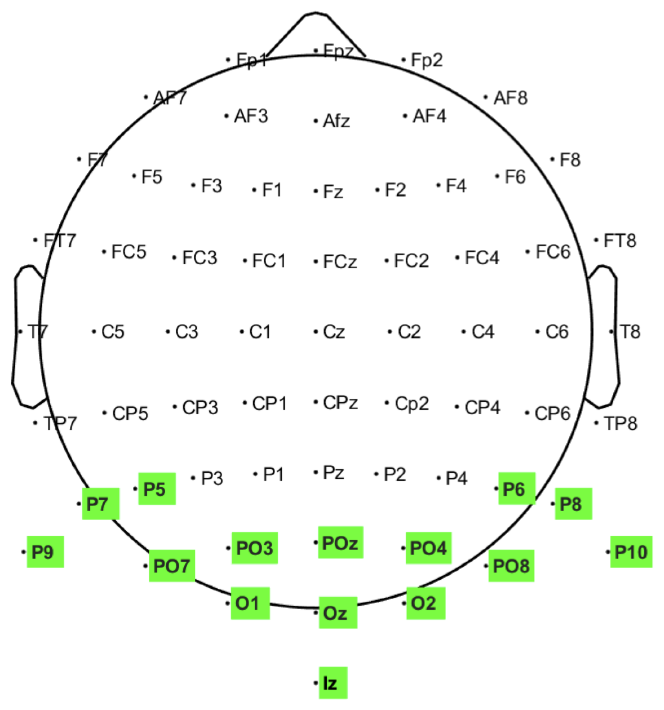
**Locations of the occipitoparietal electrodes (highlighted in green) used in our analysis**.

## 5 RESULTS

### 5.1 H1 and H2: Neural Anticipatory Intents

In this study, we sought to extend our understanding of the neural underpinnings involved in an intent-to-select interaction within a dynamic VR environment where visual information is processed over a larger area, as opposed to constrained visual fields. We assessed this using an object selection task where users intentionally selected targets, and a control condition in which they gathered information about a target (intent-to-observe interactions). We examined ERPs that were time-locked to the interaction trigger, with each trigger based on a 750ms dwell threshold.

We hypothesized **(H1)** that users in the Intent-to-select condition will anticipate a stimulus feedback from the target as a result of the selection and hence elicit an SPN, in contrast to intent-to-observe interactions. This is evident in the amplitude topographical maps shown in Figure 6, where noticeable negativity emerges from the occipital ROI in the averaged -400ms to -300ms window during intent-to-select interactions only. Our second hypothesis **(H2)** posited an SPN’s absence in the Intent-to-observe with feedback condition, as users, despite stimulus feedback, did not anticipate it due to the lack of intent to select the target. As seen, intent-to-observe interactions—both with and without feedback—displayed minimal negativity, indicating that eliciting of the SPN wave is dependent on the user’s intention.

**Figure 6:**
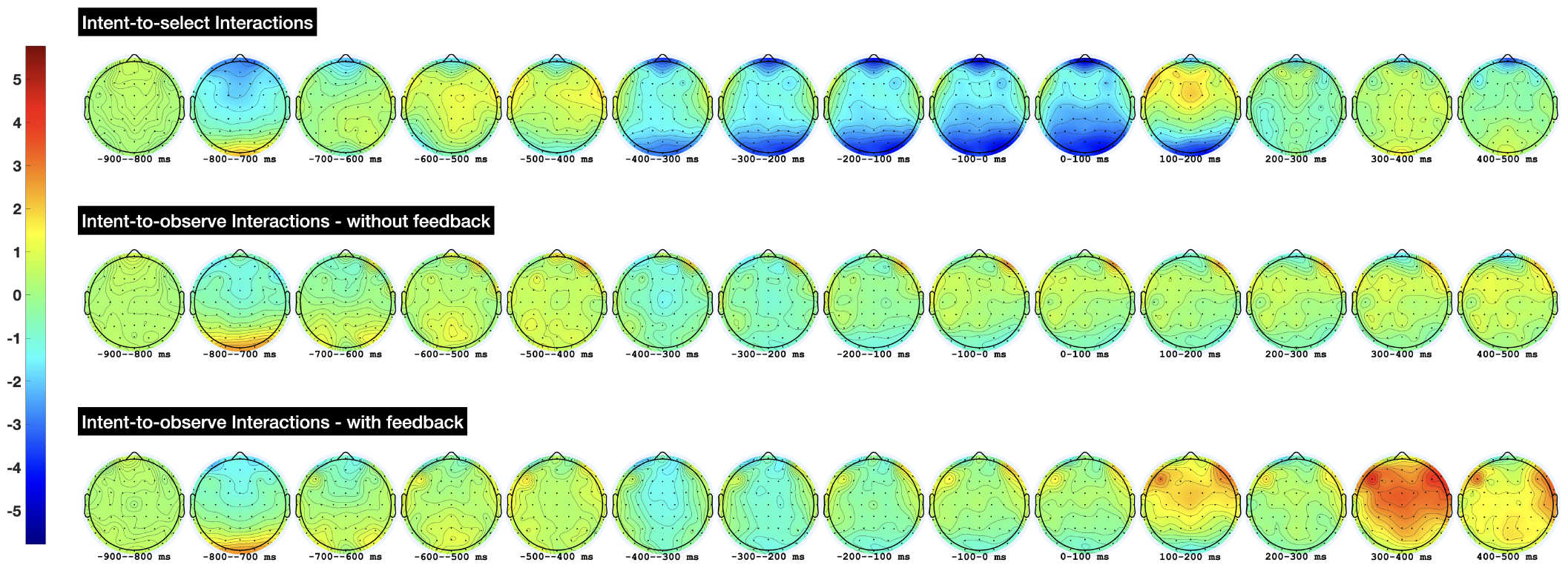
**Grand average (n = 17) topographical distributions for the mean amplitude in 100ms intervals for the period -900 to 500ms, time-locked to the interaction trigger; for the conditions: Intent-to-select (Top), Intent-to-observe without feedback (Middle), and Intent-to-observe with feedback (Bottom)**.

In Figure 7, we look more closely at the grand average (n = 17) ERPs, time-locked to the interaction trigger and averaged across the occipital ROI (O1, Oz, and O2 electrodes) because this region showed a strong emergence of negativity in the topographical maps and prior work’s focus on this region [104, 129]. The left plot contrasts Intent-to-select with the Intent-to-observe without feedback condition and the right plot contrasts Intent-to-select with the Intent-to-observe with feedback condition. For the Intent-to-select condition, the ERPs for both the ascending and descending tasks are plotted. As expected, the ERPs depict a Lambda wave positively peaking at -750ms for all three conditions. A Lambda wave is typically observed in the occipital ROI during post-saccadic eye movements, and is said to be involved in the visual processing of new information [100, 111, 125]. Another observation of interest is the Visually Evoked Potential from the target turning green, peaking at 300ms for the all but the Intent-to-observe condition where feedback was not provided (left plot).

**Figure 7:**
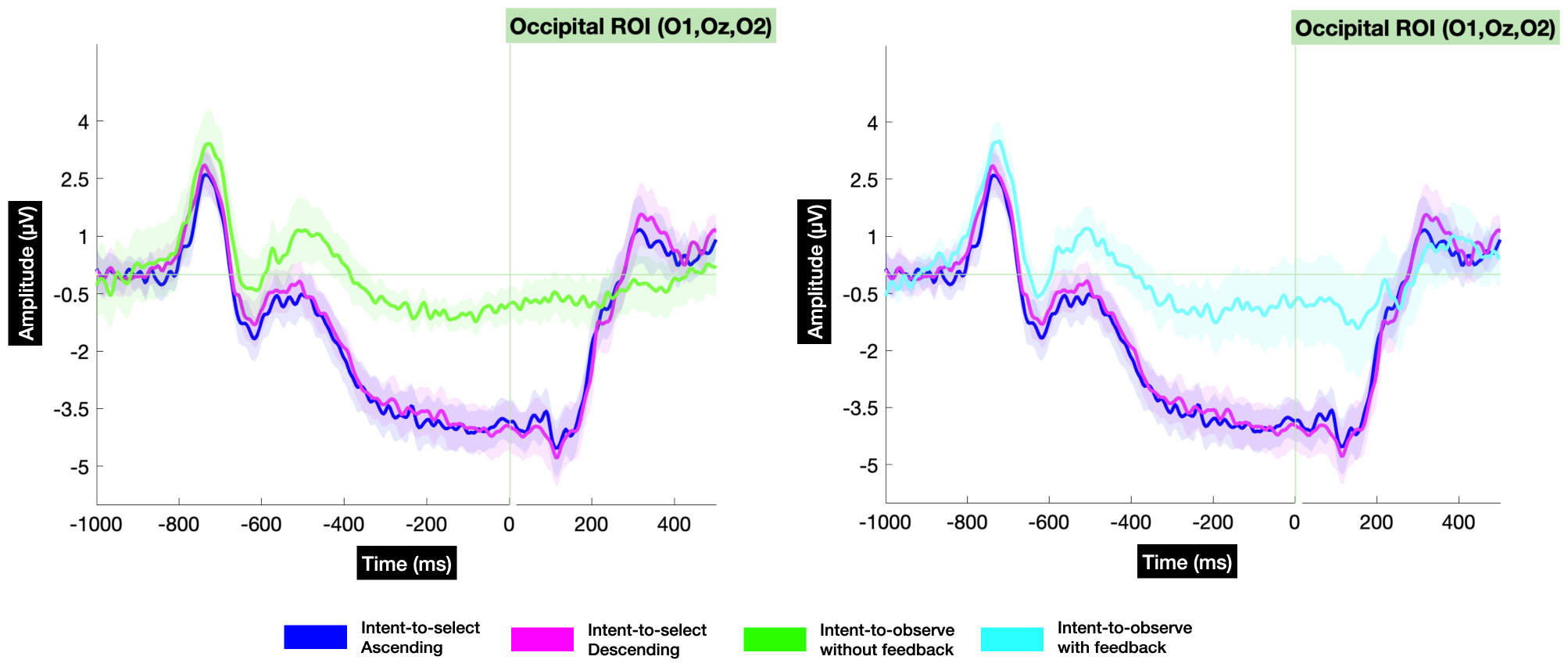
**Grand average ERPs (n = 17) at the occipital ROI (O1, Oz, O2), time-locked to the interaction trigger. The left plot contrasts Intent-to-select ERPs with Intent-to-observe without feedback ERPs. The right plot contrasts Intent-to-select ERPs with Intent-to-observe ones where feedback was provided. The shading depicts one standard error**.

#### 5.1.1 Mass Univariate Analysis

To determine statistically significant effects among the three conditions, we analyzed the ERPs via a one-way within-subjects ANOVA using a mass univariate (MU) approach from the Factorial Mass Univariate Toolbox [31]. The test included all data points from -750 to 0ms (spanning the entire dwell time duration), sampled at 256 Hz at all 15 electrodes, totaling 2895 comparisons. We did not specify a time window based on visual inspection as it can introduce notable bias [73]. Not specifying a window also allowed us to determine at what latency the intent-to-select interaction can be distinguished from an intent-to-observe one. We used a cluster-based permutation test variant based on the cluster mass statistic [9] for multiple comparisons correction, setting the family-wise alpha level at 0.05. An ANOVA was performed for each comparison using both the original data and 10,000 random within-participant permutations. For every permutation, *F* value corresponding to uncorrected *p* values of 0.05 or lower were grouped into clusters. Electrodes within 5.44 cm were seen as spatial neighbors, and adjacent time points were treated as temporal neighbors. The sum of the *F* value within each cluster defined its “mass.” The most extreme cluster mass from each of the 10001 tests was recorded and used to estimate the distribution of the null hypothesis. Clusters in the original, unpermuted data larger than 95% of clusters in the null distribution were deemed significant (*p*< 0.05).

This permutation test approach was preferred over traditional mean amplitude ANOVAs as it offered superior spatial and temporal precision while still accounting for the multitude of comparisons [31]. Furthermore, spatiotemporal averaging fails to offer specifics about the timing of the effect onsets and offsets or the precise electrodes where the effect was reliably present [36]. The choice of the cluster mass statistic was due to its proven efficacy for wide-ranging ERP effects like the P300 [36, 78], and for its efficacy in detecting clear effects from noise [36]. We used 10,000 permutations to estimate the null hypothesis distribution, as this number significantly exceeds Manly’s [77] recommendation for a family-wise alpha level of 0.05.

The results from the MU analysis are illustrated in Figure 8 with the main effect on the first raster diagram, followed by the raster diagrams of the pairwise *post-hoc* tests. Clusters with a significant effect (*p* < 0.05) are emphasized using a color gradient based on the *F* value. A notable main effect of the task condition was evident on a broadly distributed cluster that began at -750ms at PO7 and shifts posteriorly to O1 until 0ms. Other significant clusters were observed at Oz (−473ms to -242ms), Iz (−504ms to -333ms), P10 (−750ms to -629ms and -438ms to -98ms), PO8 (−453ms to 0ms), and O2 (−477ms to 0ms).

**Figure 8:**
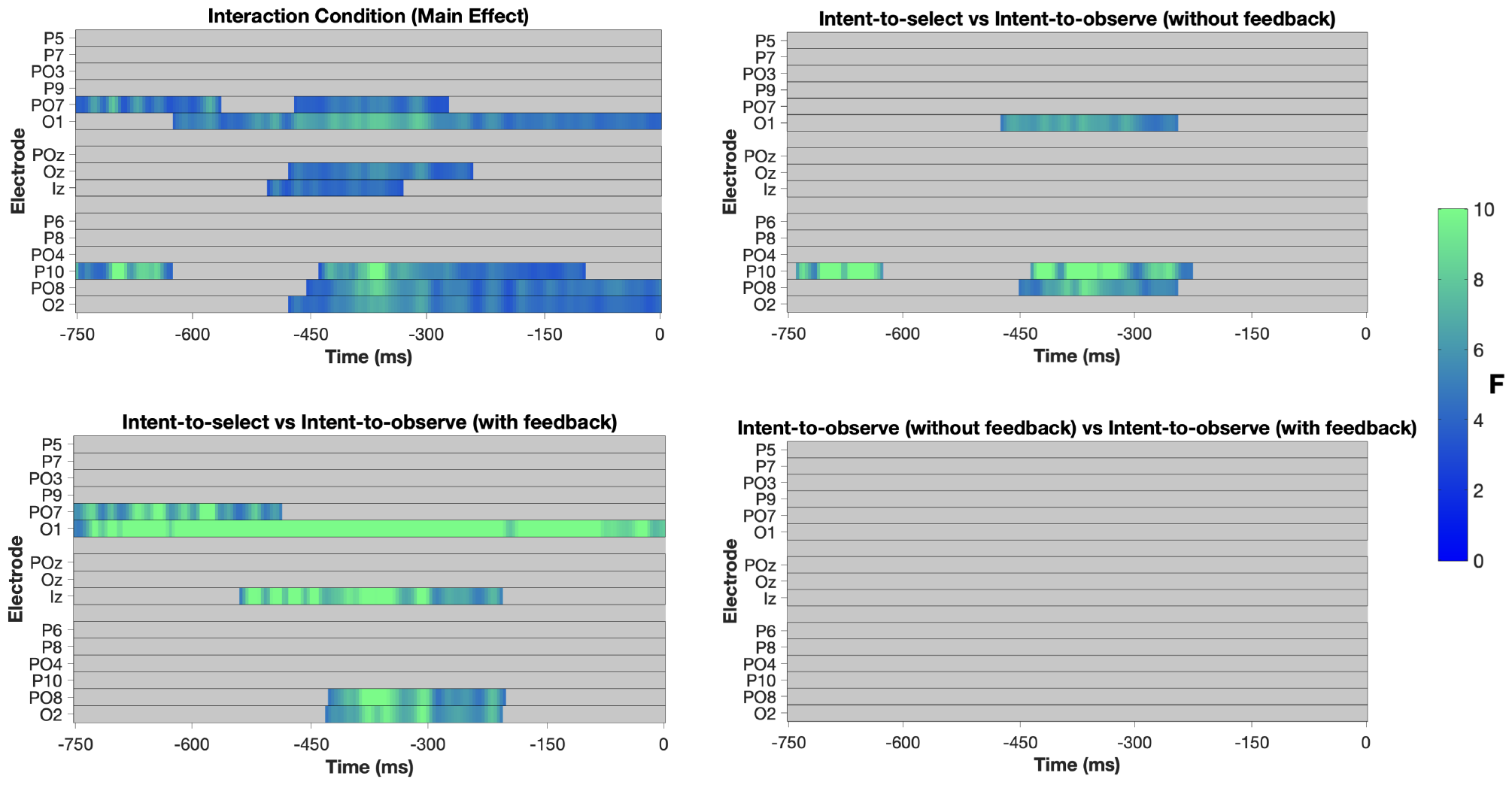
**Raster diagrams illustrating significant effects among the three conditions, based on a cluster-level mass permutation test. Note that the electrodes are arranged topographically on the y-axis (left to right hemisphere). The first diagram illustrates the main effect of the task conditions on the occipitoparietal electrodes, in the range to -750 to 0ms range, followed by the pairwise *post-hoc* tests. The colored rectangles represent the significant clusters (*p* < 0.05), with the gradient tied to the *F* value**.

For the *post-hoc* test comparing Intent-to-select versus Intent-to-observe without feedback conditions, significant clusters were identified at O1 (−473ms to -246ms), P10 (−738ms to -629ms and -434ms to -227ms), and PO8 (−449ms to -246ms). This confirms a significant difference in negativity between intent-to-select and intent-to-observe interactions for these electrodes and time points. These findings substantiate **H1** that an SPN is elicited during an intent-to-select interaction, as opposed to an intent-to-observe one in an immersive environment.

When contrasting the Intent-to-select versus Intent-to-observe with feedback conditions, a broad significant clusters was observed at O1, ranging the full duration of the dwell-threshold (−750ms to 0ms). Other significant clusters were observed at PO7 (−750ms to -488ms), Iz (−539ms to -207ms), PO8 (−426ms to -203ms), and O2 (−430ms to -207ms). These significant findings substantiate **H2**, that an SPN will be absent during intent-to-observe interactions despite feedback being presented, driving the conclusion that the SPN is influenced by the user’s intention to select the target. Lastly, no significant differences were found between the two intent-to-observe conditions - with and without feedback, further confirming that the negativity is not influenced by the presence of feedback alone.

### 5.2 H3: Target placement acclimatization

In everyday human-computer interactions, users often know the locations of frequently-used UI elements, eliminating the need to search for a desired button. Given this, we examined how familiarity with UI placements impacts an SPN during intent-to-select interactions. Our hypothesis (**H3**) posits that as users become familiar with target locations, the SPN will manifest more rapidly. We intentionally kept the target positioning consistent for every block, allowing participants to familiarize themselves with their placements. Participants went through all 5 phases (10 rounds of intent-to-select tasks, totalling 150 interactions for each participant) in a block with the same target positioning. The target positioning was randomized at the start of each block.

We hypothesized that as participants learned target placements, the time to complete the intent-to-select tasks would reduce phase-by-phase, as a result of reduced target searches. This trend is evident in the left plot of Figure 9, which shows the mean task completion times for the five phases, averaged across participants and blocks. An ANOVA revealed a main effect (*F* (4,80) = 7.2, *p* < 0.001), with Tukey’s *post-hoc* tests confirming a significant difference between phase 1 and phase 5 times at *p* < 0.001. Accordingly, we compared the grand averaged (n = 17) ERPs in phase 1 (1471 trials with 1.9% rejected) with phase 5 (1429 trials with 2.8% rejected). The middle plot of Figure 9 illustrates the topographical distributions for the -400 to -300ms time window, time-locked to the interaction trigger. The top and bottom rows depict phase 1 and phase 5, respectively. Using a 0.1 Hz high-pass filter (HPF), as detailed in Section 4.6, a marked increase in negativity is observed for phase 5 compared to phase 1, suggesting a faster emergence of the SPN. However, a one-tailed within-subject MU *t*-test based on the cluster-level mass permutation approach did not reveal any significant effects (*p* > 0.05) for the difference wave (phase 1 ERP - phase 5 ERP). 2500 random permutations were tested and the test included all data points from -750 to 0ms, sampled at 256 Hz at all 15 electrodes, totaling 2895 comparisons. We used the MU ERP Toolbox [36], and our choice for MU analysis remains the same as stated in Section 5.1.1.

**Figure 9:**
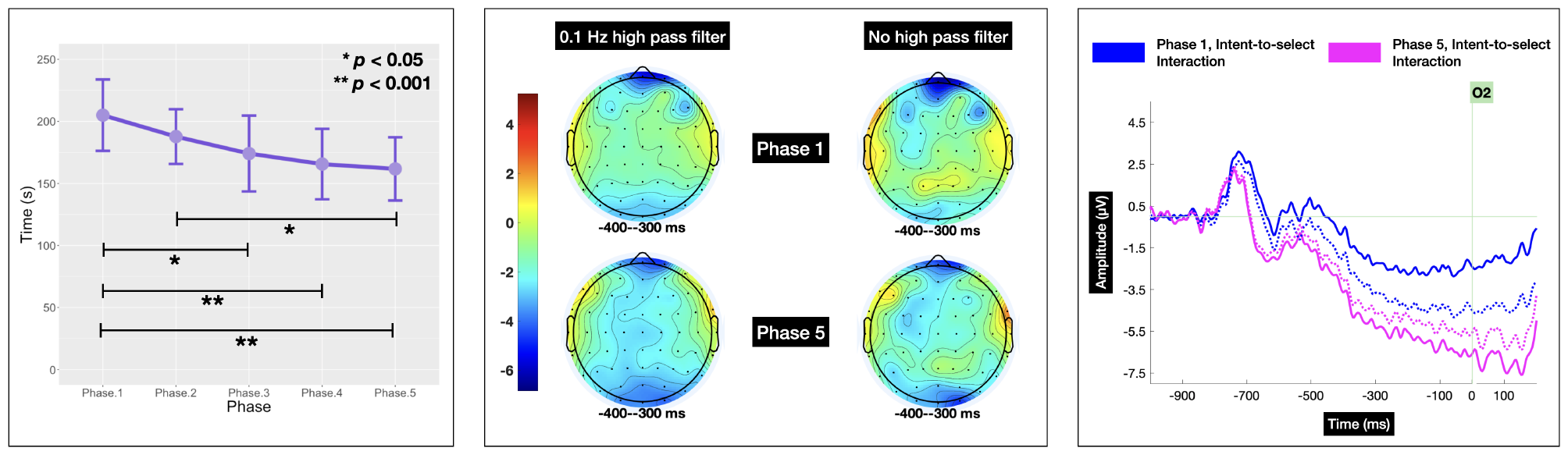
**(Left) Mean task completion time per phase for the Intent-to-select condition, averaged across blocks and participants; error bars represent one standard deviation. (Middle) Grand average (n = 17) mean amplitude distributions for phases 1 and 5 averaged across blocks, with and without a high-pass filter. (Right) Grand average (n = 17) ERPs from O2 for phases 1 and 5 averaged across blocks. Dotted lines show ERPs with a 0.1 Hz high-pass filter, while solid lines show ERPs without it**.

In line with previous works that examined EEG negativity in intent-to-select versus intent-to-observe interactions [104, 129], we omitted the HPF. Interestingly, in Figure 9’s scalp maps, without the HPF, a more prominent emergence of negativity is seen for phase 5 than phase 1 during the same time window, specifically at O2. Figure 9’s right plot displays the grand average (n = 17) ERPs for O2 in phases 1 and 5, averaged across all blocks. Dotted lines depict ERPs with the 0.1 Hz HPF, while solid lines represent ERPs without it. We see a considerably larger difference in the ERPs without the HPF, in contrast to when it’s applied. We used the fractional area latency metric [53, 69] to compare onset timings between phases 1 and 5, identifying the latency (onset) at which 50% of the SPN’s negative AUC has been reached. With the HPF, phase 5’s SPN covered 50% of the negative AUC 23.44ms earlier, with its latency measured at -226.56ms, compared to phase 1’s latency of -203.12ms. Without the HPF, phase 5’s SPN was 35.15ms sooner, with its latency measured at -226.56ms, compared to phase 1’s -191.41ms.

Notably, when the HPF is removed, we see a significant cluster for the difference wave at O2 starting from -516ms until 0ms, as seen in the bottom plot of Figure 10, with the color gradient depicting the *t* score. The parameters for the test was similar as for the one with a HPF. The top plot of Figure 10 illustrates the mean amplitude distribution (top) and the *t* score distribution (bottom) for the difference wave with and without the HPF. Interestingly, only O2 shows a significant effect without the HPF, which could be attributable to the fact that an SPN is stronger in the right hemisphere [6].

**Figure 10:**
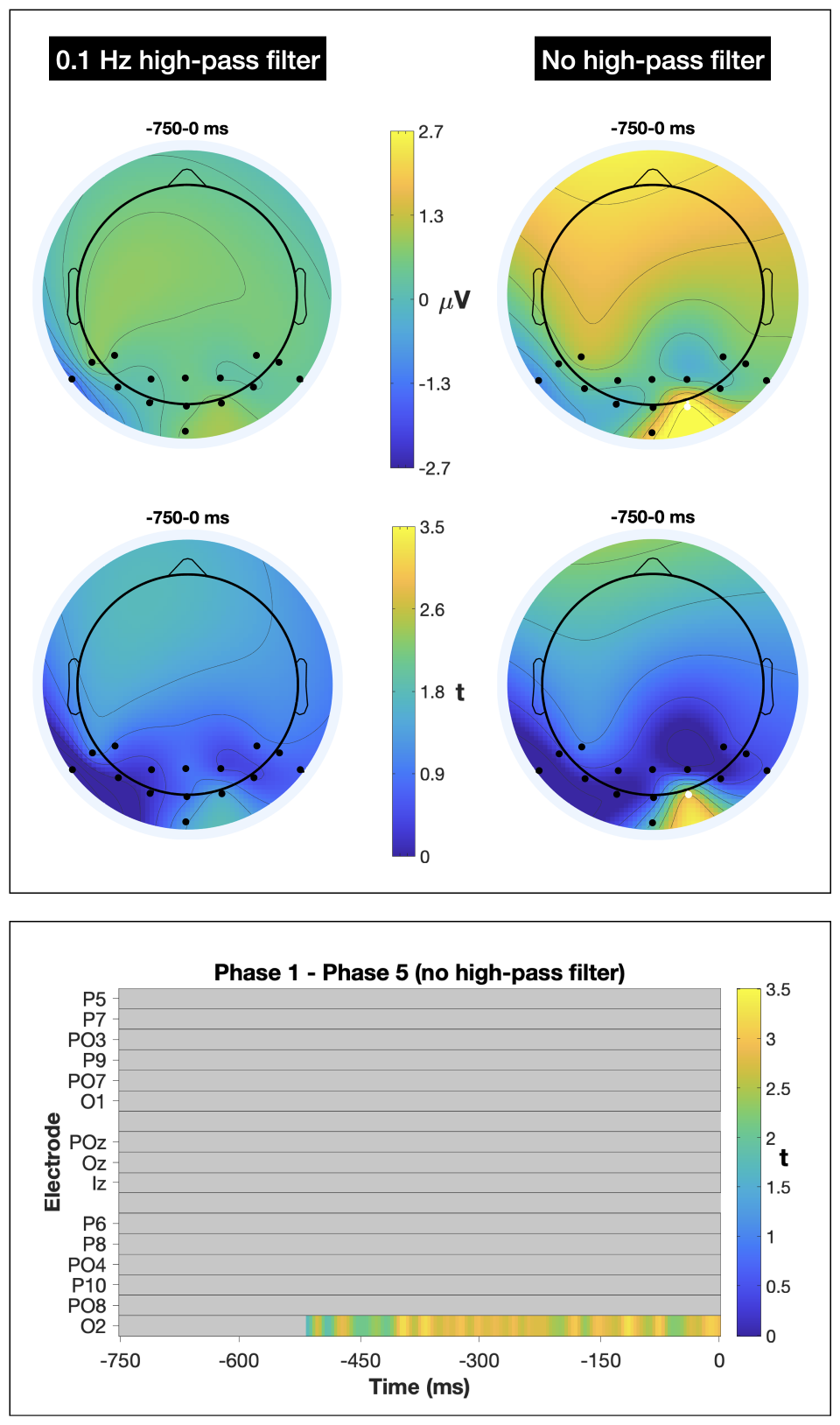
**(Top) Mean topographies of the phase 1 - phase 5 difference wave with amplitude distribution on the top and *t* score distribution on the bottom, for both with and without a high-pass filter. O2 was significant in the no filter condition. (Bottom) Raster diagram illustrating significant differences between phase 1 and phase 5 ERPs without a high-pass filter, based on a cluster-level mass permutation test. Colored rectangles indicate time points/electrodes at which a significant effect was observed, with the gradient tied to the *t* score**.

We would like to emphasize that a HPF is highly advised to to mitigate low-frequency environmental noise, including artifacts from factors such as user perspiration-induced slow drifts [48, 72].

This is particularly crucial for real-world BCIs employing active dry electrodes with high impedances, even though it may influence slow ERP components [79]. Consequently, we retained this filter for **H1** and **H2**. The absence of statistically significant differences in the ERPs with the HPF makes it challenging to reject our null hypothesis for **H3**. Nevertheless, the significance without the filter further strengthens our direction that the SPN emerges more rapidly when users become familiar with the target placement. The implications of this finding can assist real-time BCI systems to optimize the time windows used for classification depending on the user’s familiarity with the visually focused target.

## 6 CONTRIBUTIONS, BENEFITS, AND LIMITATIONS

The Midas touch dilemma has long plagued gaze-based interactions. Researchers have addressed the issue in a myriad ways (see Section 2.1), mostly involving an extra selection step requiring conscious physical effort. A natural and passive approach to compliment the intuitiveness of gaze-based pointing is lacking. Our study delved into the potential of leveraging the SPN as a passive indicator of user intent. We observe higher negativity levels during intent-to-select interactions, peaking at -4μV at 0ms or when the target was selected. In contrast, the control conditions exhibited minimal negativity with levels approaching -1μV, aligning with findings by Shishkin et al. [104]. Interestingly, the intent-to-select ERPs (Figure 7) bear close resemblance to those observed in Zhao et al. [129]’s work.

Furthermore, prior work [104, 129] did not investigate the lateral parietal ROI (P5, P6, P7, P8, P9, and P10 electrodes). Notably, in our research, the P10 electrode yielded particularly significant results (Figure 8). Pronounced negativity for intent-to-select interactions is also evident at the frontal ROI (FpZ in Figure 6). While not investigated here, we attribute this to anticipation of the audio cue on successful selection, given that frontal areas display greater negativity for auditory stimuli and occipital areas for visual ones [7, 60].

Our results also show robust statistical significance via the MU approach, which not only corrects for multiple comparisons but also harnesses the full spatial and temporal precision of EEG; unlike prior work [104, 129] that employed a mean amplitude ANOVA. Additionally, the MU approach highlights the lower bounds of the temporal onset or the earliest moment the EEG signal responds to a forthcoming stimuli [88]. These insights are crucial for BCI development, indicating how soon the brain can distinguish between conditions. This, in turn, suggests the potential rapidity of a BCI system in differentiating intent-to-select from intent-to-observe interactions. In our study, Intent-to-select versus Intent-to-observe without feedback interactions differed as early as 12ms post initial gaze at P10. For the Intent-to-select versus Intent-to-observe with feedback scenario, this distinction was evident at both PO7 and O1 from -750ms itself (when the user looked at the target).

We demonstrated that an SPN can differentiate between intent-to-select and intent-to-observe interactions within an immersive VR context. This is particularly noteworthy given the intricacies of immersive environments, characterized by frequent head movements, diverse depth of fields, and expansive field of views. Despite the fact that we use a VR environment specifically, we believe our results are generalizable to the broader XR umbrella as many of the concepts related to the three-dimensional aspect remain similar.

Additionally, our research elucidates that SPN is intrinsically tied to a user’s intention to select the target and is not influenced by feedback alone — a vital confounding element overlooked in earlier studies [104, 129]. The implications of this finding bolster the rationale for the SPN wave’s elicitation, and reinforces its practicality for a BCI system. This finding can also be interpreted from prior literature on SPN [11] which states that an SPN is present only for feedback that conveys certain information. In our research, this information could simply be a notifying the user of a successful selection, provided they were actively aiming to select it. Moreover, these insights pave the way for BCI designs that dynamically adjust feedback in response to real-time neural activity, creating a biofeedback mechanism. For instance, as a user focuses on a UI element, the system assesses negativity levels and responsively adjusts the stimulus to heighten the anticipatory response, possibly facilitating faster selection, similar to Crispin et al. [13]. If the user did not intend to select the target, this adaptive stimulus shift will not impact the negativity.

In regard to **H3**, we were unable to show a statistically significant difference with a HPF. This might stem from participants not being entirely accustomed to the target placements even by the fifth phase. We did not objectively measure participants’ level of familiarity, but instead inferred from their task completion times. Moreover, we did not instruct participants to familiarize themselves with the target locations. We do, however, see a strong significant difference at O2 without a HPF, strengthening the direction of our hypothesis. Given the importance of a HPF in a BCI, we believe additional research and a targeted approach is necessary to validate our third hypothesis. Interestingly, in phase 5 ERP without a HPF (right plot of Figure 9), we also observe a heightened negativity exceeding -6μV at 0ms. This suggests that practice and the entrenched memory of the feedback can foster stronger negativity, aligning with previous findings that practice enhances intentional regulation in slow-cortical potentials [83].

The approach we adopted in this work emphasizes the implicit and fluent nature of these interactions and decouples BCI paradigms from the constraints of the information transfer rate [101], by which we not only streamline the system’s functionality but also broaden its applicability. This ensures that the system is not just an assistive tool for disabled users but becomes an inclusive interface that healthy users can adopt. An added advantage is that the system could be further streamlined using only a few electrodes in the occipitoparietal ROI. For instance, a HMD might incorporate 3 to 5 electrodes on its backstrap, resulting in a more user-friendly form factor. The integration of eye-tracking with implicit neural activity thus not only emerges as a pivotal direction in fostering wider acceptance and integration of BCIs in everyday scenarios, but it also presents a more unobtrusive approach to addressing the Midas touch dilemma.

### 6.1 Limitations and Future Work

We have identified limitations that future research can address. Firstly, the Dwell technique can lead to user fatigue [37, 42, 98]. While prior work have tried to identify an optimal threshold duration, a definitive one remains unestablished [75, 98, 116]. Second, we did not develop a classification model, and hence cannot compare how well (accuracy) an intent-to-select interaction can be distinguished from an intent-to-observe one for single trial ERPs. Furthermore, as this study is an initial exploration of the SPN’s potential in tackling the Midas touch problem, it is not yet feasible to compare our approach with an established technique that resolves this issue. Yet, our robust statistical findings hint at the potential to design a hBCI leveraging the SPN, and has the capability to be a faster, effortless, and fluent interaction paradigm than SSVEPs or P300-based BCIs.

Third, further research on real-time preprocessing, ocular artifact correction, and addressing strong motion artifacts, is vital to effectively mainstream this anticipatory interaction paradigm for real-time use. Gherman and Zander [35] found that different postures did not have an impact on a passive BCI’s classifier performance, strengthening our case for a real-time system. Fourth, it should also be noted that our high-resolution EEG data originated from a controlled lab environment using research-grade EEG equipment. Exploring the efficacy of this approach in real-world scenarios with dry electrodes, warrants further investigation. Fifth, intent-to-select and intent-to-observe interactions were restricted to their own conditions, further research is necessary in examining these neural signatures when these conditions are intermixed, like they would be in a real world setting.

Finally, our study did not include a condition decoupling feedback from intent-to-select interactions to ascertain the feedback’s role in SPN elicitation. However, prior work has demonstrated the SPN’s absence without feedback [11] and the necessity of temporal predictability or instilled memory of the target providing feedback for SPN emergence [8, 21, 22]. This aligns with our findings in Figure 9’s right plot and as stated earlier, phase 5 ERP shows greater negativity than phase 1 ERP, due to a more instilled memory of the feedback by phase 5. Furthermore, the SPN’s prominence in the occipital region, known for processing visual stimuli, confirms that it is linked to the visual feedback [8]. Future research can examine the SPN’s behavior in the sudden cessation of feedback during intent-to-select interactions.

#### 6.1.1 Future Work

Our next efforts are to build a model to establish the approach’s accuracy in offline classification, before advancing to real-time classification. Correcting ocular artifacts for real-time online classification presents a challenge as ICA-based removal is a time-consuming process, though prior work [44] has effectively implemented it with negligible time delays. We plan to explore additional corrective methods like SGEYESUB [59], which remove ocular artifacts in real-time after an initial calibration phase to record specific eye artifacts for building a correction model. Moreover, examining the behavior of the SPN wave in the absence of ocular artifact corrections, will provide insights regarding the necessity of these corrections in an online framework. Other preprocessing steps, including resampling, re-referencing, line noise elimination, filtering, and baseline correction (applied post-detection of a *gaze on target* event), do not pose a time constraint issue for real-time implementation.

We hypothesize that an online implementation could be improved by fine-tuning the model to a user’s specific neural signature, requiring a prior calibration step involving Dwell-based selections. It’s imperative that the model attains a high accuracy to prevent user frustration and guarantee smooth interactions; we plan to evaluate this using subjective and qualitative metrics. To counteract false negatives, an online system should incorporate a fail-safe confirmation mechanism, such as a pinch gesture or a voice interface. On the other hand, false positives can be addressed by implicitly detecting user states such as ErrPs (discussed in Section 2.3). We also plan to integrate natural gaze dynamics, such as differentiating between ambient and focal fixations [61], which could enhance classification accuracy and optimize computational resources.

In light of evolving interaction paradigms, future work may also explore more sophisticated techniques, such as object manipulation or adjusting sliders, moving beyond mere selection tasks to enrich user experience. A promising direction would be linking the SPN to a variable dynamic value, as opposed to the binary classification employed in selection tasks. Lastly, while our method holds tremendous potential, it’s crucial to investigate privacy-preserving techniques, especially because these systems can decode user states without their explicit awareness, potentially leading to unintended retrievals of sensitive cognitive information.

## 7 CONCLUSION

In this research, we explored the use of neural activity as a confirmation trigger to address the Midas touch dilemma in gaze-based interactions. Specifically, we empirically examined anticipatory EEG signals to serve as a passive mental switch, signifying a user’s intent to select a visually-focused object within an XR setting. Our results confirmed two primary hypotheses: intent-to-select interactions elicit an SPN wave, differentiating them from intent-to-observe interactions, and the elicitation of the SPN wave hinges on user intent. Furthermore, while initial evidence suggests that familiarity with target placement might lead to a quicker emergence of the SPN, this was not conclusively established. Our findings underscore an SPN’s capability to discern between intent-to-select and intent-to-observe gaze-based interactions, paving the way for more effortless, passive, and efficient BCIs, particularly for XR environments.

## Footnotes

1 https://www.hp.com/us-en/vr/reverb-g2-vr-headset-omnicept-edition.html

2 https://www.biosemi.com/products.htm

3 https://unity.com

4 https://github.com/sccn/labstreaminglayer

